# DNA Methylation (DM) data format and DMtools for efficient DNA methylation data storage and analysis

**DOI:** 10.1101/2023.11.27.568779

**Authors:** Qiangwei Zhou, Cong Zhou, Zhixian Zhu, Yuanhui Sun, Guoliang Li

## Abstract

DNA methylation is a critical epigenetic mechanism that plays a pivotal role in various biological processes. Currently, larger datasets from whole-genome bisulfite sequencing for DNA methylation pose challenges throughout the computational analysis pipeline, including storage and memory constraints. Unfortunately, storage formats and analysis tools have not kept pace with these increased resource demands. In this study, we present a new and efficient design for storing DNA methylation (DM) data after mapping in compressed binary indexed DM format. Our format significantly reduces storage space by 80%-95% compared to commonly used file formats for DNA methylation data after mapping. To enhance the processing of DNA methylation data in DM format, we have developed DMtools, a comprehensive toolkit that offers utilities such as rapid and random access, computation of DNA methylation profiles across genes, and analysis of differential DNA methylation. The analysis speed is improved by over 100 times compared to existing methods. Furthermore, we have created pyDMtools, a Python package that efficiently processes DM format files for Python users. The integration of the DM format and its associated tools represents significant progress in handling and exploring DNA methylation data, offering the potential to significantly reduce storage needs and improve downstream analysis capabilities.

## Introduction

DNA methylation is a critical epigenetic mechanism that plays a pivotal role in various biological processes, such as cell differentiation (1,2), development (3), aging (4), genomic imprinting (5), X chromosome inactivation (6) and other important biological processes (7). Aberrant DNA methylation can lead to tumor formation (8,9). Whole Genome Bisulfite Sequencing (WGBS) is a powerful method for precisely detecting DNA methylation at a single-base resolution across the entire genome. It achieves this by converting nonmethylated cytosines into thymines, significantly advancing the study of DNA methylation (10).

Currently, widely-utilized tools in DNA methylation data analysis include BatMeth2, Bismark, BS-Seeker, and BSMAP (11-14). Although efficient in handling DNA methylation data alignment and methylation level calculations, these tools store output files in various text formats. This storage format poses challenges in terms of extensive storage space, slow downstream analysis speed, and difficulty in extracting specific methylation levels for a chromosome region or gene, as well as computing methylation profiles for particular genes. This problem becomes more marked when dealing with multiple-sample data, leading to prolonged analysis times and increased storage pressure. For example, the National Center for Biotechnology Information (NCBI) already possesses over 5500 sets of WGBS data, and there are also several DNA methylation databases such as ASMdb (15) and MethBank (16) that include thousands of bisulfite sequencing (BS-Seq) datasets, imposing substantial computational and storage burdens. Therefore, there is an urgent need for a universal compressed binary index format and a downstream analysis tool that can support all DNA methylation alignment software, minimize storage space requirements, and accelerate downstream DNA methylation data analysis.

In the realm of DNA methylation data analysis, our introduced DNA Methylation (DM) data format and the accompanying DMtools tackle pivotal challenges related to file size, downstream analysis speed, and functionality. Notably, DMtools is a pioneering manipulation tool tailored specifically for the binary-indexed DNA methylation data format. The DM format files produced by our approach demonstrate a significantly reduced size, approximately 5%-20% compared to commonly used DNA methylation formats. While recent tools like CGmaptools(17) and BAllCools(18) have proposed CGbz and BAllC compression formats, respectively, it is noteworthy that CGmaptools still relies on downstream analysis provided by text formats, and BAllCools lacks downstream analysis functionality. In contrast, our work achieves superior compression ratios, accelerates downstream computation by over a hundredfold, and provides a comprehensive solution for efficient storage and analysis of DNA methylation data.

### DM format and DMtools

The DM format is designed for efficient storage and swift querying of DNA methylation data. It encompasses DNA methylation details, and indexing information, as illustrated in Figure 1A. This format effectively stores genome-wide DNA methylation levels at single base/region resolution, providing comprehensive information (detailed in Table 1). The inclusion of genome location indexing enables rapid retrieval of DNA methylation results for any genomic region. These features ensure quick localization of target regions with less time complexity, streamlining DNA methylation queries through expeditious data parsing and access.

**Table 1.**
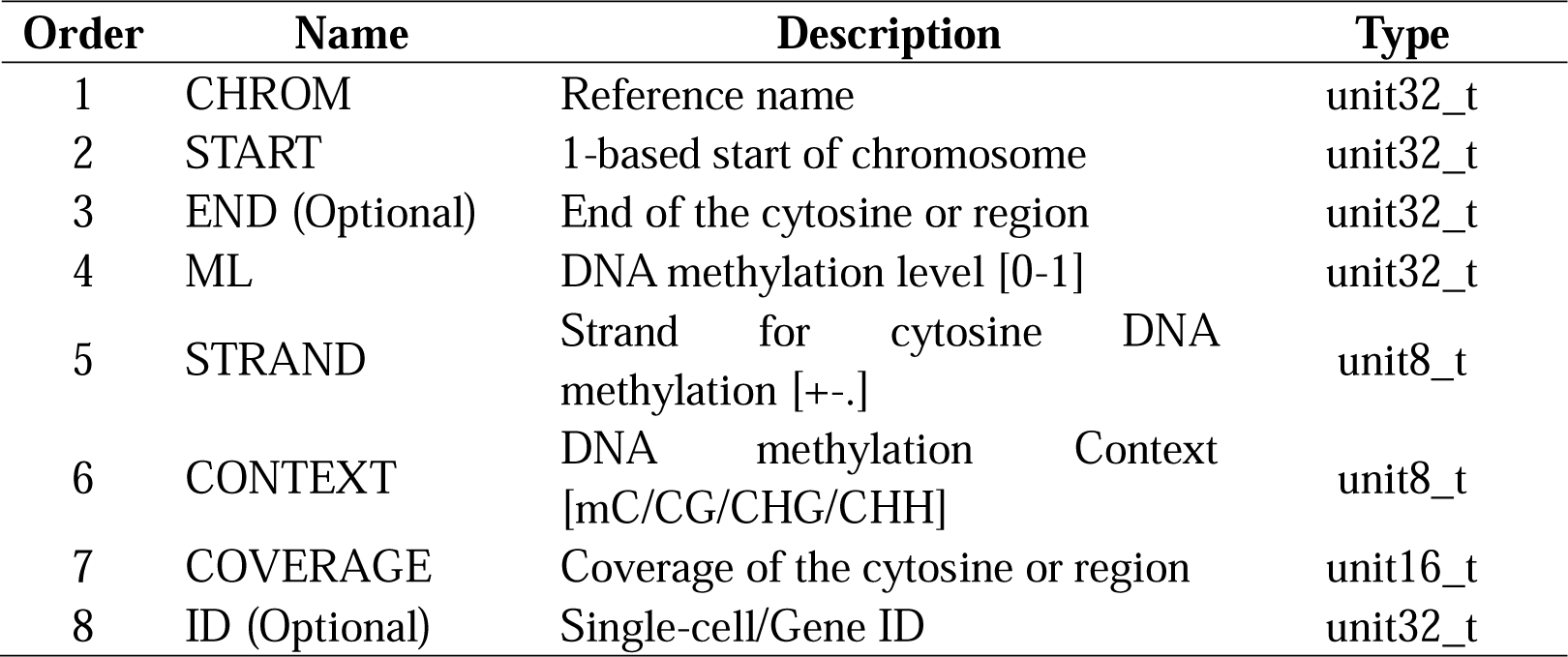
Basic information in DM format and DMtools.

| Order | Name | Description | Type |
| --- | --- | --- | --- |
| 1 | CHROM | Reference name | unit32_t |
| 2 | START | 1-based start of chromosome | unit32_t |
| 3 | END (Optional) | End of the cytosine or region | unit32_t |
| 4 | ML | DNA methylation level [0-1] | unit32_t |
| 5 | STRAND | Strand for cytosine DNA methylation [+-.] | unit8_t |
| 6 | CONTEXT | DNA methylation Context [mC/CG/CHG/CHH] | unit8_t |
| 7 | COVERAGE | Coverage of the cytosine or region | unit16_t |
| 8 | ID (Optional) | Single-cell/Gene ID | unit32_t |

**Fig. 1.**
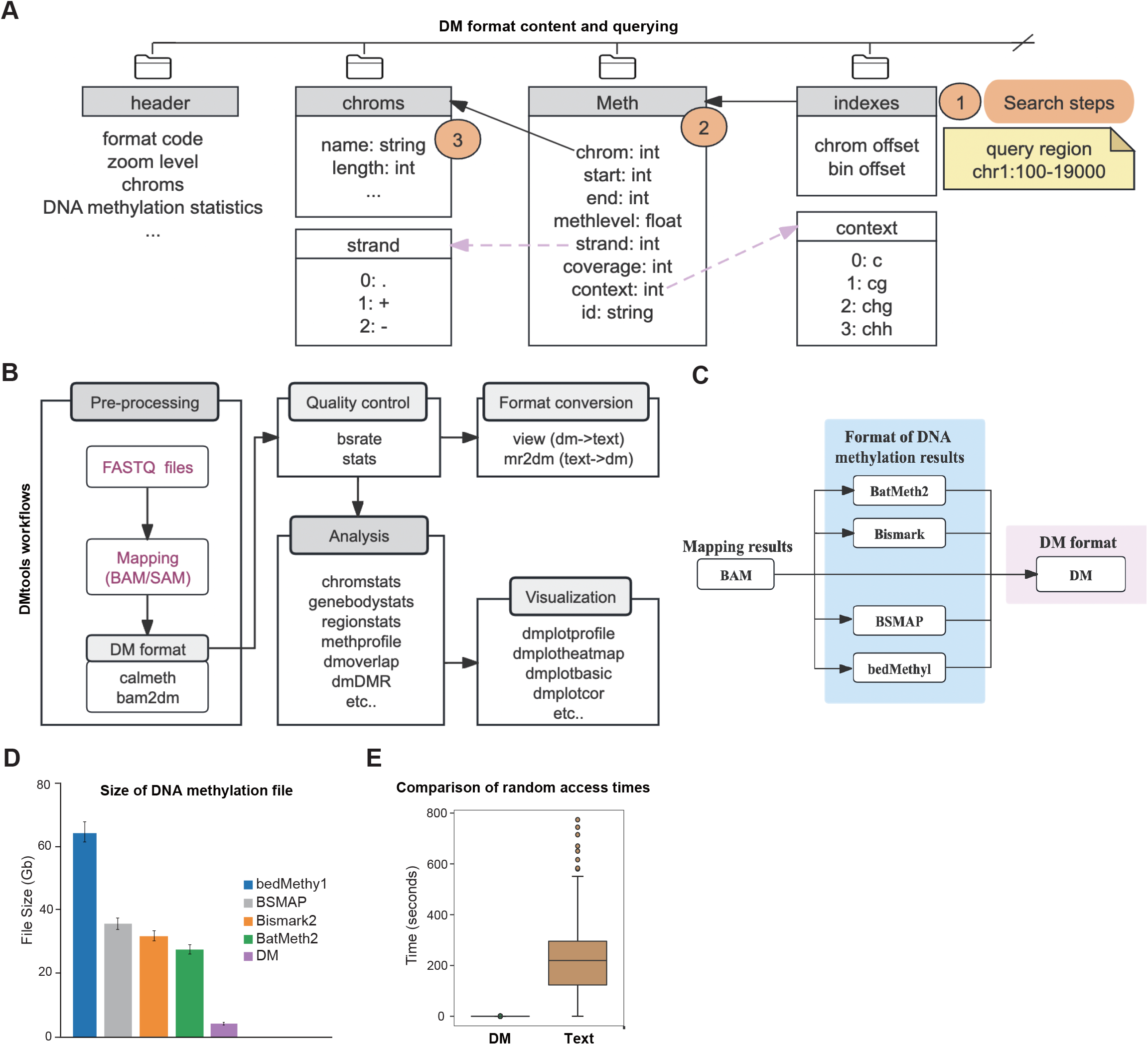
DM format and related performance comparison. **(A)** Detailed information of the DM format and an example of querying in a random region. Step 1) Represents the search query region. Step 2) Searches and locates the query region based on the DM file index. Step 3) Retrieves and returns DNA methylation result information within the query region. **(B)** DMtools workflows and tools. Entry points for external data are highlighted in purple. **(C)** Conversion from other formats of DNA methylation files after mapping to DM format. **(D)** Comparison of storage space for different DNA methylation file formats. File formats are DM-format and tab-separated text DNA methylation results from BatMeth2, Bismark, BSMAP, and bedMethyl format. **(E)** Comparison of analysis speed for chromosome region DNA methylation level calculation with DM and tab-separated methylation file formats.

We have further developed DMtools, a powerful C++ tool designed to facilitate the manipulation of DNA methylation files in the DM format (Figure 1B). It offers seamless operations on DM files and a comprehensive range of downstream analysis functions. DMtools can convert data from popular analysis software like BatMeth2, Bismark, BSMAP and bedMethyl into the DM format (Figure 1C), ensuring compatibility and flexibility. DMtools also offers a comprehensive range of downstream analysis functions. These include the assessment of bisulfite conversion rate, rapid calculation of DNA methylation levels for genomic regions and gene methylation profiles, as well as robust differential methylation analysis (Figure 1B).

Compared to traditional tab-delimited text file formats for DNA methylation data, such as BatMeth2 format, Bismark format, BSMAP format and bedMethyl, the DM format takes up less storage space (reduces about 80-95%) (Figure 1D and Supplementary Fig. S1). For querying and computing in any specific region, as shown in Figure 1A, DM format determine the precise location of the queried DNA methylation information in the file based on the index (Step 1). It then retrieves the detailed DNA methylation information (Steps 2-3) and calculates the DNA methylation results for the region. With the DM format, query and computation speed can be significantly accelerated. Notably, the utilization of DMtools for computing regional DNA methylation levels and gene profiles has achieved a remarkable speed improvement of over a hundred-fold (Figure 1E).

Except for its excellent analytical capabilities, DMtools offers a variety of DNA methylation calculation and visualization features. These include the sample correlation analysis (Supplementary Fig. S2, S3), calculation of coverage information (Supplementary Fig. S4), categorization of DNA methylation level (Supplementary Fig. S5), generation of DNA methylation profiles upstream and downstream of genes or transcription start sites (Supplementary Fig. S6-S8), and the generation of heatmaps displaying DNA methylation for all genes (Supplementary Fig. S9-S11). Moreover, for differential DNA methylation analysis (Supplementary Fig. S12), DMtools require only about 10% of the memory compared to methylKit (19), making it a reliable and efficient choice for DNA methylation research (Supplementary Fig. S13-S16). To facilitate the efficient handling of DM format files by users, we also provide a Python package called pyDMtools.

### DMtools software and datasets used for demonstration

DMtools is a C++ package that utilize the htslib and libBigWig libraries to parse and manipulate DNA methylation levels in the DM format. With DMtools, users can convert sorted BAM files of read alignments, as well as other DNA methylation formats like Bismark or bedMethyl methylation formats, into DM format. Additionally, we have developed a Python package, pyDMtools, based on DMtools that enables users to read and process DM files in Python. Detailed instructions on how to use DMtools and the pyDMtools Python package for reading and processing DM files are readily available on the DMtools website (https://dmtools-docs.readthedocs.io/en/latest/index.html). Installing pyDMtools is a straightforward process using either ‘pip3 install pyDMtools’ or ‘conda install -c bioconda pyDMtools’.

We demonstrated the capabilities of DMtools with genome-wide DNA methylation data results for human NeuN control, NeuN Schizo, and AZA-AML3 (20) to assess the file size, differential methylation analysis, and visualization. Additionally, we obtained WGBS-Seq data for Arabidopsis Col-0, met1-3, ddcc, and mddcc (21), and used DMtools to analyze and visualize the DNA methylation levels. Please refer to the supplementary file for details of RRBS and target-BS data.

## Conclusion

With the application of next-generation sequencing technologies and their decreasing costs, more and more WGBS and RRBS DNA methylation data are becoming available. Efficient storage and processing pipelines are necessary to handle such large datasets. While existing methods such as GZIP, CGbz and BAllC are available for compressing DNA methylation results, they lack suitable tools for subsequent analysis of DNA methylation data. Additionally, GZIP is not well-suited for effective compression of DNA methylation results (22).

In this work, we have presented an innovative binary index DM format for DNA methylation data after mapping, which includes methylation level, coverage, context, and additional relevant information. Compared to traditional DNA methylation formats, the DM format offers several advantages, such as smaller file sizes and efficient query capabilities for DNA methylation levels in any genomic region. We have developed DMtools, a comprehensive toolkit that can generate DM format files from sorted BAM files, convert traditional DNA methylation files to DM format, process DM format files, and perform various DNA methylation analyses, including DNA methylation calculation, differential methylation analysis, and DNA methylation visualization.

We believe that the DM format and DMtools have the potential to make the analysis of WGBS and RRBS DNA methylation data more efficient. We are committed to continuously improving the software and incorporating new features based on user feedback in future.

## Data availability

The software and usage documents can be accessed via the following links:

Software documentation: https://dmtools-docs.rtfd.io/

GitHub repository for DMtools: https://github.com/ZhouQiangwei/DMtools

GitHub repository for pyDMtools: https://github.com/ZhouQiangwei/pyDMtools

## Acknowledgements

We thank the group members for providing feedback on the software.

