## Supplemental Materials for "DNA Methylation (DM) data format and DMtools for efficient DNA methylation data storage and analysis"

Zhou et al., 2024

DMtools is a software suite designed for the analysis and visualization of DNA methylation data. It supports a variety of DNA methylation data types, including whole-genome bisulfite sequencing (WGBS), reduced representation bisulfite sequencing (RRBS), targeted bisulfite sequencing, and more. The software can be used for quality control, differential methylation analysis, correlation analysis, visualization, and other tasks related to DNA methylation data analysis.

In terms of data analysis, DMtools can perform quality control on sequencing data by filtering low-quality reads and removing adapter sequences. It can also map sequencing reads to a reference genome and calculate DNA methylation levels for each cytosine position. Differential methylation analysis can be performed to identify regions of the genome with significant changes in methylation between different samples or groups. Correlation analysis can be performed to explore relationships between different samples or groups.

DMtools also includes various visualization tools to help users interpret and present their data. These tools include heatmaps, scatterplots, and boxplots to visualize DNA methylation levels and coverage across the genome. Additionally, DMtools can generate track files compatible with popular genome browsers like UCSC and IGV, making it easy to visualize DNA methylation data in the context of other genomic features.

Overall, DMtools is a powerful tool for the analysis and visualization of DNA methylation data, making it useful for researchers in fields such as epigenetics, genomics, and bioinformatics.

### Evaluation of bisulfite conversion rate

For bisulfite sequencing data, the level of DNA methylation is estimated by converting unmethylated cytosine (C) to thymine (T) and then calculating the percentage of T at sites where the reference genome is C. The accuracy of this estimation depends on the bisulfite conversion rate, which directly affects the percentage of C to T. To evaluate the conversion rate of bisulfite, lambda sequences are typically added during library construction. The Lambda sequence is a completely unmethylated sequence, so after bisulfite treatment, all C sites should be converted to T with 100% efficiency.

However, some WGBS data do not include lambda sequences for estimating the conversion rate of bisulfite. In such cases, we can use several alternative methods to evaluate the conversion rate: (1) calculating the mitochondrial methylation level (applicable to some mammals); (2) calculating the DNA methylation level of CHH (applicable to some animals); (3) calculating the methylation level of a specific chromosome specified by the user; and (4) calculating the methylation level of chloroplasts (applicable to some plants). It is important to note that some studies have reported DNA methylation in mitochondria, so using mitochondrial DNA methylation level to estimate the conversion rate of bisulfite requires careful consideration.

### DNA methylation level calculation

DMtools provides an 'regionstats' command that can be used to calculate the DNA methylation level within a user-specified region of interest. This functionality is highly useful for researchers who may be interested in the methylation level of specific genes, transposable elements (TEs), or other regions of interest. By using the dmtools regionstats command, researchers can quickly and easily calculate the methylation level of each region of interest, and perform further analysis and comparison. This is crucial for uncovering the role of DNA methylation in various biological processes and its association with diseases.

To calculate the methylation density level in a given genomic region, only cytosines with coverage greater than the preset threshold are used. The DNA methylation level in a genomic region is defined as the total number of sequenced Cs over the total number of sequenced Cs and Ts at all cytosine positions across the region, and the equation is as follows:

$$ML(x)_{method} = \begin{cases} \dfrac{\Sigma_{i=1}^{N} C}{\Sigma_{i=1}^{N}(C+T)}, & \text{if method = weighted} \\ \dfrac{\Sigma_{i=1}^{N} ML(x_i)}{N}, & \text{if method = mean} \end{cases}$$

where N is the total number of cytosine sites whose coverage is more than the predefined threshold in the genomic region.

**Datasets used in DMtools for illustration**

1. human AML3 WGBS-Seq data (2 replicates for wild type and 2 for treated).

   AZA-treated: GSM1329865, GSM1329866

   AZA-WT: GSM1329867, GSM1329868

   This dataset is used for partial visualization and differential analysis. DNA methylation inhibitors of DNA methyltransferases (DNMT) can reverse the characteristic abnormal promoter DNA methylation and related gene silencing in specific acute myeloid leukemia (AML) patients (Lund, et al., 2014).
2. Arabidopsis WGBS-Seq data.

   GEO ID: GSE183987, GSE169497

   WT, met1, ddcc and mddcc: wild type and different mutants

   This test data is used for partial visualization. Arabidopsis DNA methylation data includes whole-genome DNA methylation data of wild-type Col-0, met1 mutant, ddcc mutant, and mddcc mutant. The data information, with NCBI GEO ID, is GSE183987 and GSE169497 (Zhao, et al., 2022).
3. RRBS.

   GEO ID: GSM3603286, GSM3603287, GSM3603288

   This test data is used for validating the DM file format storage and DMtools' basic functionality. (Pastore, et al., 2019)
4. Targeted-BS.

GEO ID: GSM659626, GSM659627, GSM659628, GSM659629

This test data is used for validating the DM file format storage and DMtools' basic functionality. (Lee, et al., 2011)

In the following sections, we utilized DM format and DMtools to analyze DNA methylation data from various selected samples. We employed multiple datasets including WGBS, RRBS, and target-BS to validate the versatility of the DM format and the multifunctionality of DMtools. Furthermore, based on the DM format, we conducted separate tests for various DNA methylation level calculations and visual analyses. Finally, we presented differential DNA methylation analysis tests using DMtools.

### 1. Analysis process and data description

In this example, we applied DM format and DMtools to analyze DNA methylation data from DNMT variants and wild type.

After downloading the WGBS-Seq data, quality filtering was performed, and DNA methylation sequence alignment was obtained using BatMeth2 (Zhou, et al., 2019) to obtain a SAM file. The SAMtools (Li, et al., 2009) tool was then used to filter low-quality reads (mapQ >= 30) and sort and convert them into BAM format. Subsequently, DMtools were used to calculate DNA methylation levels, data coverage, and other information.

### 2. Calculating DNA methylation levels from BAM files and writing to DM format

First, DNA methylation levels are calculated based on the sorted BAM format. The script is as follows: dmtools bam2dm -g hg38.fa -b $sample.sort.bam -m $sample.dm --zl 0 -C -S --Cx. Additionally, if a bw file is needed for visualization in the IGV browser, DMtools can be used to generate a file containing only methylation levels and with a zoom level, as follows: dmtools bam2dm -g hg38.fa -b ${sample}.sort.bam -m ${sample}.zm2.bw --zl 2 -E.

### 3. DM format is applicable to RRBS and target-BS data.

We tested the application of DM format and DMtools in RRBS and target-BS data and compared the file sizes. The results are shown in the Fig. S1.

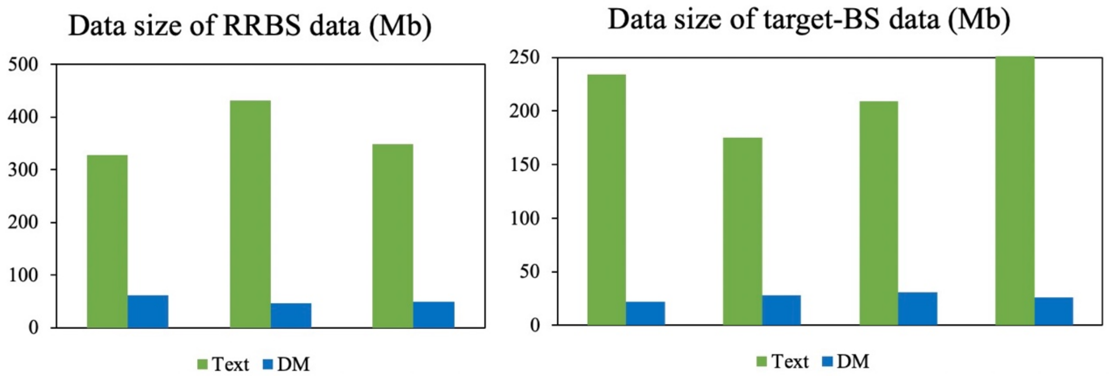


**Fig. S1. Data size for RRBS and target-BS data**

**4. Quality control analysis based on DM format.**

In DNA methylation data analysis, calculating the correlation between biological replicates and different samples is essential. Biological replicates refer to data obtained by sequencing or performing the same experiment multiple times on the same sample. By calculating biological replicates, we can evaluate the reproducibility and stability of the data, and determine whether changes in methylation levels are due to true biological differences or technical noise.

On the other hand, the correlation between different samples can help us evaluate their similarity or differences. By calculating the correlation between samples, we can identify which samples have similar methylation patterns and which samples exhibit differences. This helps us identify critical differential methylation sites and determine the similarity and differences in methylation patterns among different samples, leading to more accurate conclusions.

Therefore, in DNA methylation data analysis, calculating biological replicates and the correlation between different samples is a crucial step that helps us evaluate the reliability and consistency of the data.

We can use DMtools chromstats to calculate the DNA methylation level in genomic regions of different lengths and calculate the correlation between different samples based on the results (Fig. S2, S3).

```
python3 dmPlotCor.py --plotFile bs.correlation.pdf -m spearman -p scatterplot \
```

```
--plotNumbers -f sample1.chrmeth.txt sample2.chrmeth.txt sample3.chrmeth.txt \
sample4.chrmeth.txt -c CG -l s1_rep1 s1_rep2 s2_rep1 s2_rep2 \
--xRange 0 1 --yRange 0 1
```

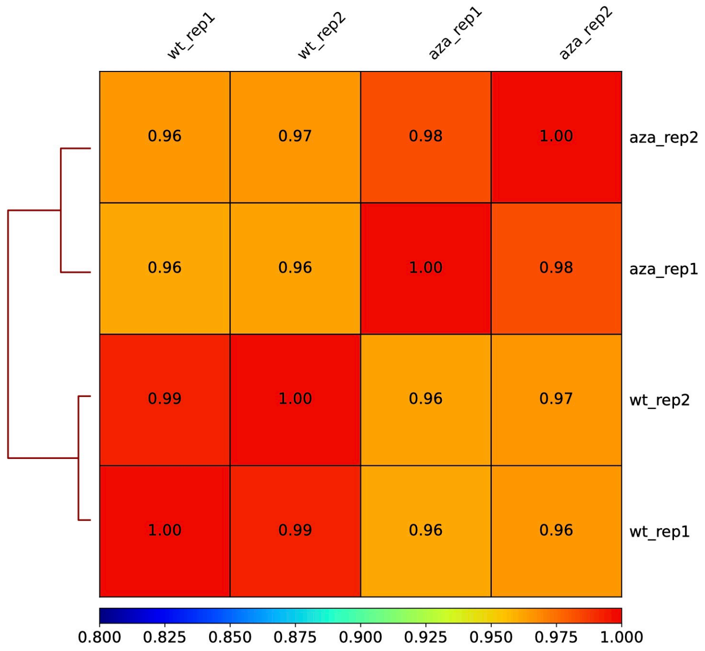


**Fig. S2. Correlation heatmap of DNA methylation samples**

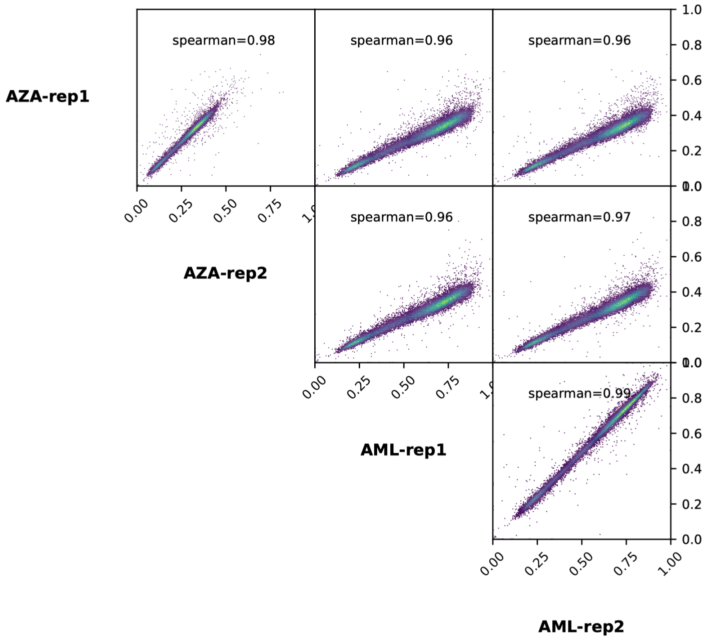


**Fig. S3. Correlation scatterplot of DNA methylation samples**

We used DMtools to calculate the DNA methylation levels of each sample and stored them in DM format. Furthermore, we calculated the coverage of cytosine sites in each sample using DMtools: dmtools stats -i ${sample}.zm0.dm -o ${sample}.cover --tc. Based on this, we used the DMtools visualization script to generate coverage statistics: python3 dmtools/bt2basicplot.py -s GSM1329865.cover.stats GSM1329866.cover.stats GSM1329867.cover.stats GSM1329868.cover.stats -o dnmt-aml.cover.pdf -l AZA-treated-1 AZA-treated-2 WT-1 WT-2. The coverage information is shown in Fig. S4. Meanwhile, we can also view the differences in DNA methylation levels among different samples, as shown in Fig. S5. We can clearly see that the high methylation sites (0.8,1] in the two mutants were significantly lower than those in the wild type. This is because the treatment with AZA affected the function of DNA methyltransferases, resulting in a significant decrease in DNA methylation levels.

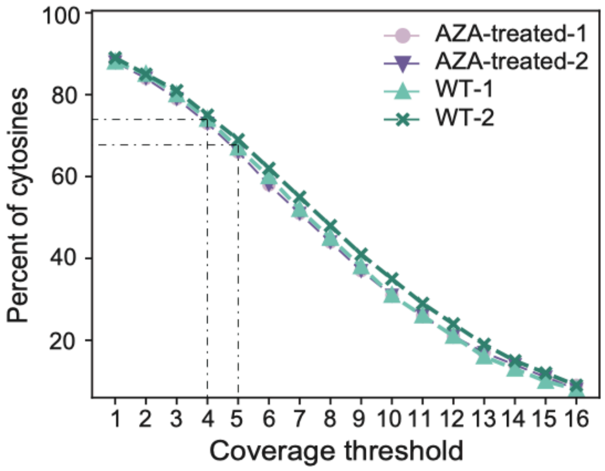


**Fig. S4. Percent of cytosines with coverage equal to or larger than the threshold in different samples**

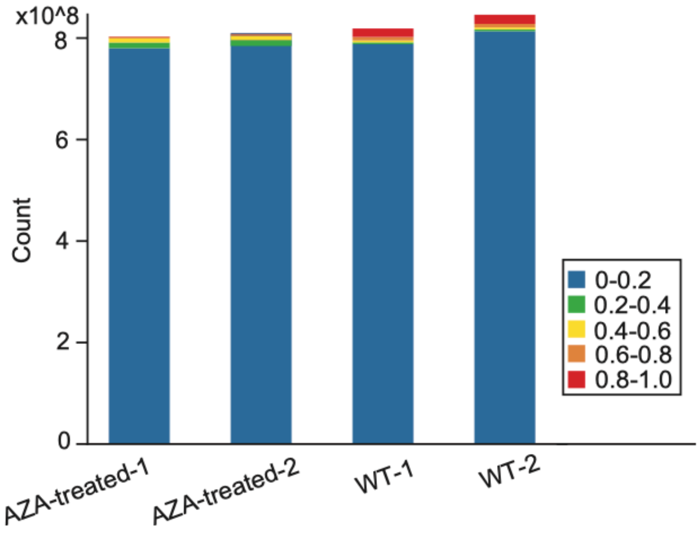

**Fig. S5. Distribution of cytosines with diverse DNA methylation levels in different samples**

**5. DNA methylation comparison analysis for two or more samples.**

DMtools provides a method to calculate the correlation analysis between two groups of samples based on DM format. Because the samples are AML-WT and AZA-treated AML data, and the DNA methylation has undergone global and dramatic changes, it can be observed that the correlation of methylation is relatively low.

In addition, DMtools provides the ability to calculate DNA methylation levels and profiles for genes or any region of interest based on the DM format. As an example, we calculate the distribution of DNA methylation levels upstream and downstream of genes and at the gene start site. The specific process is as follows: dmtools profile -i ${sample}.dm --gtf gene.gtf -o ${sample}.profile for gene body and dmtools profile -i ${sample}.dm --gtf gene.gtf -o ${sample}.profile.tss --profilemode 1 for gene start site. Based on these results, we complete the visualization work using python3 dmtools/bt2profile.py -f GSM1329865.profile.aver GSM1329866.profile.aver GSM1329867.profile.aver GSM1329868.profile.aver -o dnmt.profile.pdf -l AZA-treated-1 AZA-treated-2 WT-1 WT-2 for gene body and python3 dmtools/bt2profile.py -f GSM1329865.profile.tss.aver GSM1329866.profile.tss.aver GSM1329867.profile.tss.aver GSM1329868.profile.tss.aver -o dnmt.tss.profile.pdf -l AZA-treated-1 AZA-treated-2 WT-1 WT-2 -xl up2K TSS down2K -s 1 1 for gene start site. The visualization results are shown in Fig. S6, where we can observe the difference in DNA methylation distribution between the wild type and DNMT mutant on genes.

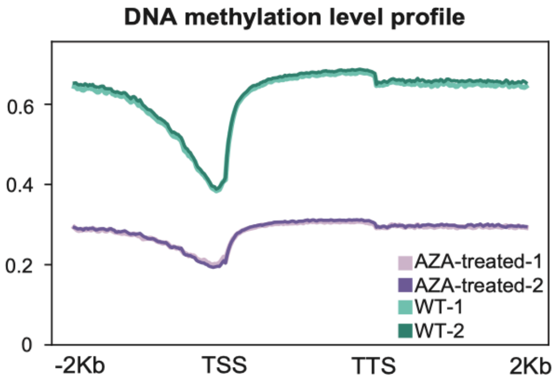

**Fig. S6. Average profiles of DNA methylation levels around genes in different samples**

Distribution patterns of mC, CG, CHG, and CHH DNA methylation levels upstream and downstream of gene transcription start sites (TSS) in different samples in Arabidopsis (Fig. S7).

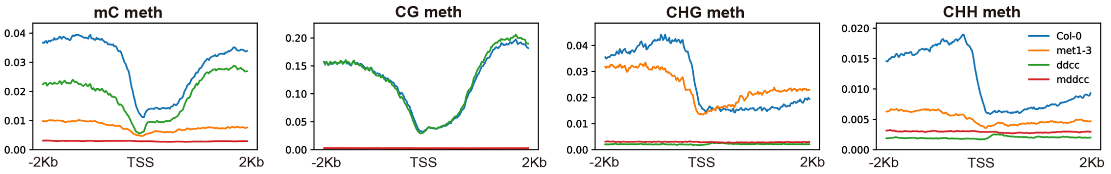


**Fig. S7. DNA methylation profile across gene TSS**

DMtools not only can calculate the distribution of DNA methylation upstream and downstream of gene TSS, but also can calculate the differential DNA methylation of CG, CHG, and CHH DNA methylation in the entire gene upstream and downstream in different samples (Fig. S8).

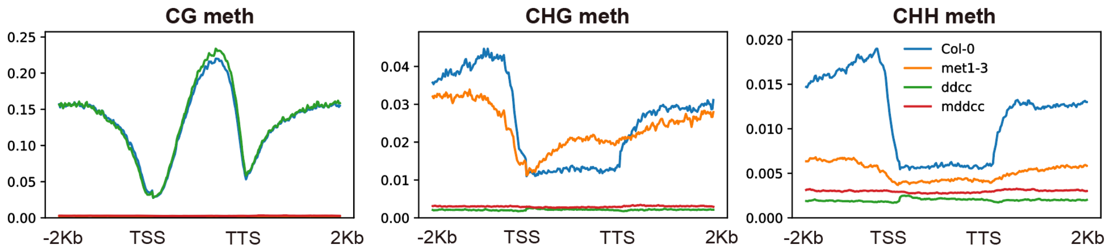


**Fig. S8. DNA methylation profile (CG/CHG/CHH) across genes**

While calculating the distribution pattern of DNA methylation on genes, we also calculated the DNA methylation results after dividing each gene into bins, and based on this result, we completed the heatmap display. Specifically:

python3 dmtools/bt2heatmap.py -m GSM1329865.profile.cg GSM1329866.profile.cg GSM1329867.profile.cg GSM1329868.profile.cg -l AZA-treated-1 AZA-treated-2 WT-1 WT-2 -o dnmt.heatmap.pdf -sl TSS -el TTS --zMax 0.8 --colorMap Spectral_r --kmeans 3. The visualization result is shown in Fig. S9.

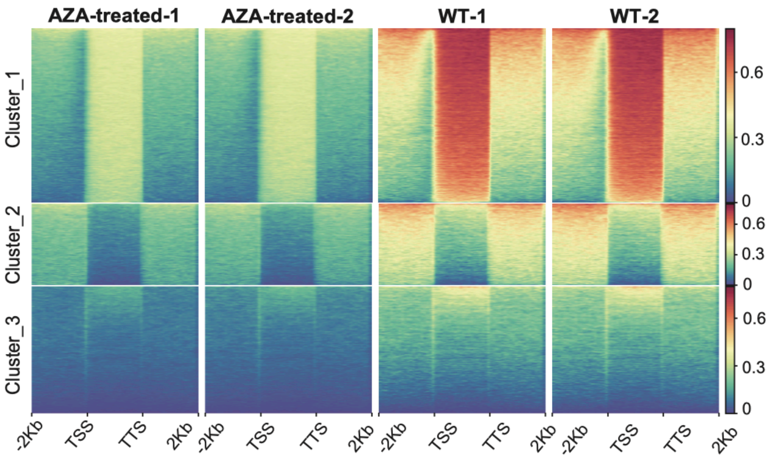


**Fig. S9. Heatmap of DNA methylation levels around all genes**

Similarly, we also performed analysis on DNA methylation data of several groups in Arabidopsis species, including WT, met1, ddcc, and mddcc, using DMtools (Fig. S10).

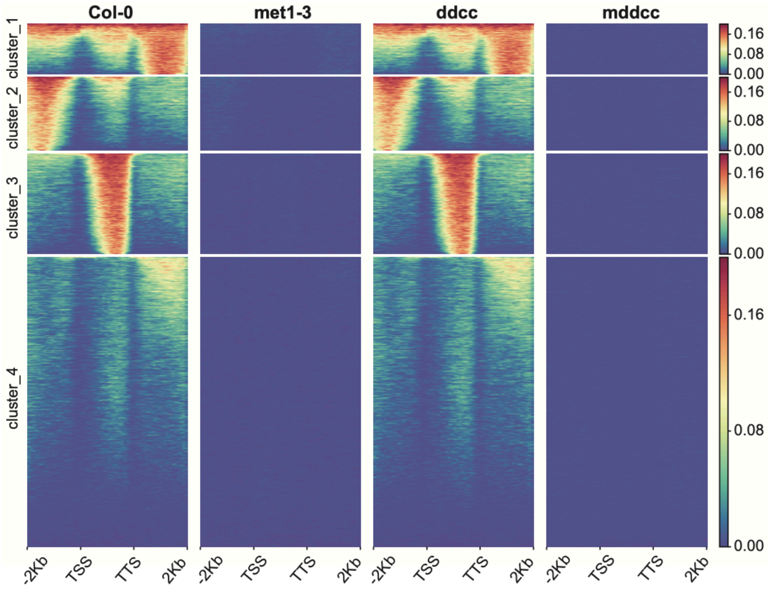


**Fig. S10. Heatmap of DNA methylation levels around all genes of different samples in Arabidopsis**

The previous section showed the differences in DNA methylation distribution patterns upstream and downstream of genes in different samples. DMtools can also quickly calculate the DNA methylation level of each gene. The specific process is dmtools bodystats -i ${sample}.bm --gff gene.gff -o ${sample}.body --printcoverage 1. After obtaining the DNA methylation levels of each gene and promoter region, heat map visualization and box plot visualization can be completed using DMtools. Heat map visualization is performed using dmtools/bt2heatmap.py, and box plot visualization is performed using dmtools/bt2basicplot.py (Fig. S11). They will not be further elaborated here.

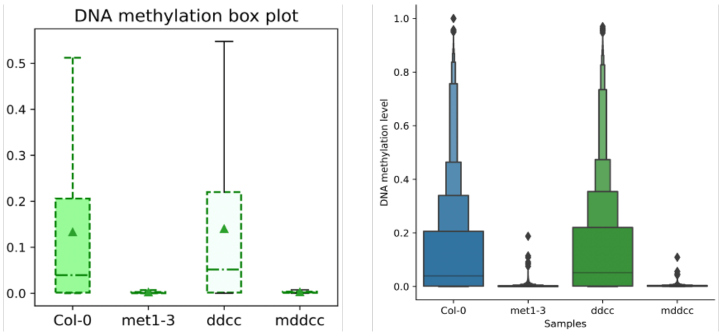


**Fig. S11. Gene methylation boxplot of different samples in Arabidopsis**

### 6. DNA methylation differential analysis

All differential analyses were performed on the same computing node in single-threaded mode. Memory usage and runtime were measured using Linux commands.

The above content shows the DNA methylation level calculation, interval DNA methylation, gene DNA methylation calculation, heatmap, and profile visualization based on the DM format using DMtools. DMtools also provides differential DNA methylation analysis algorithms based on the DM format. The specific calculation process is dmtools/dmDMR -1 GSM1329867.zm0.dm,GSM1329868.zm0.dm -2 GSM1329865.zm0.dm,GSM1329866.zm0.dm -p wt-aza.dm --mindmc 5 --minstep 200. The result file will contain differentially methylated DNA sites and differentially methylated DNA regions (Fig. S12). For differentially methylated DNA regions, the heatmap calculation and display mentioned earlier can also be used.

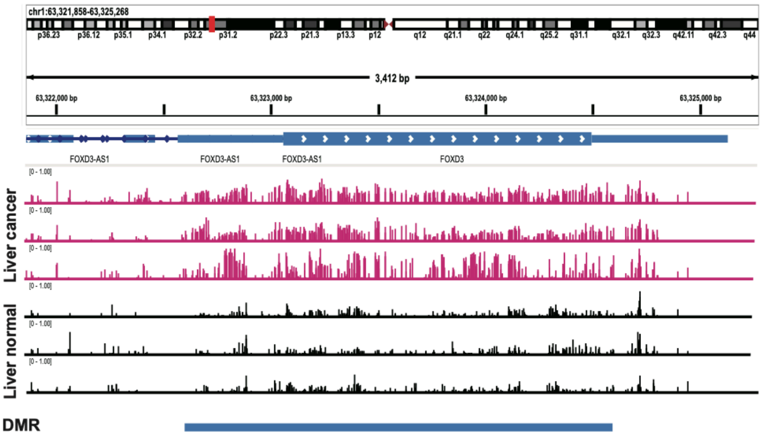


**Fig. S12. Screenshot of DNA methylation levels and DMR with IGV**

We compared the memory usage, running time, and differential DNA methylation results of DMtools and methylKit (Akalin, et al., 2012) for differential analysis.

6.1 Memory usage statistics for different software.

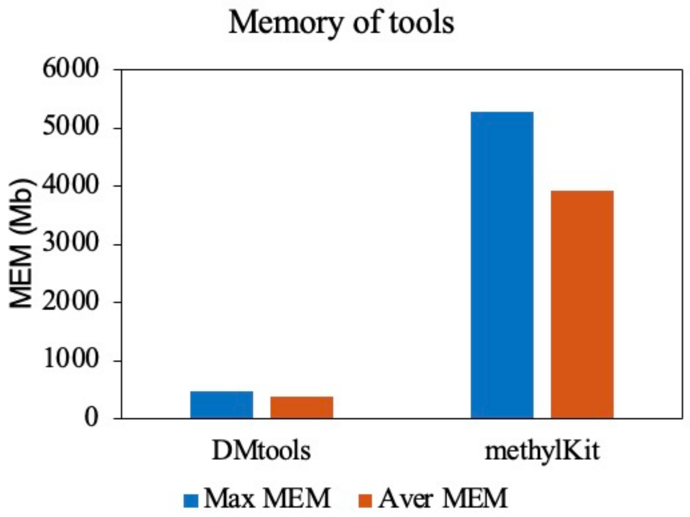


**Fig. S13. Memory required for differential analysis.**

6.2 Running time statistics for different software.

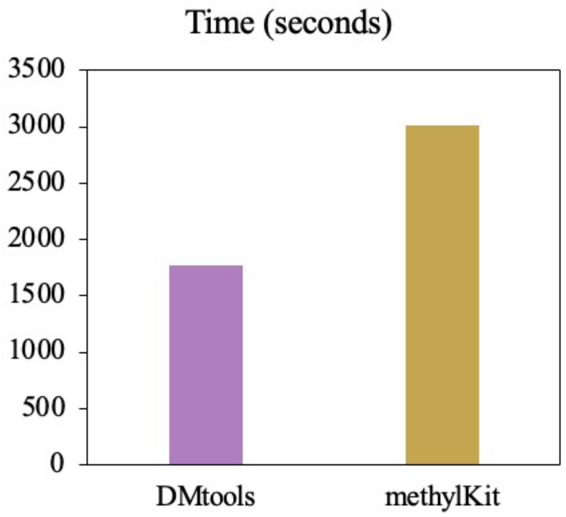


**Fig. S14. Time used for differential analysis.**

6.3 Overlap of differential DNA methylation

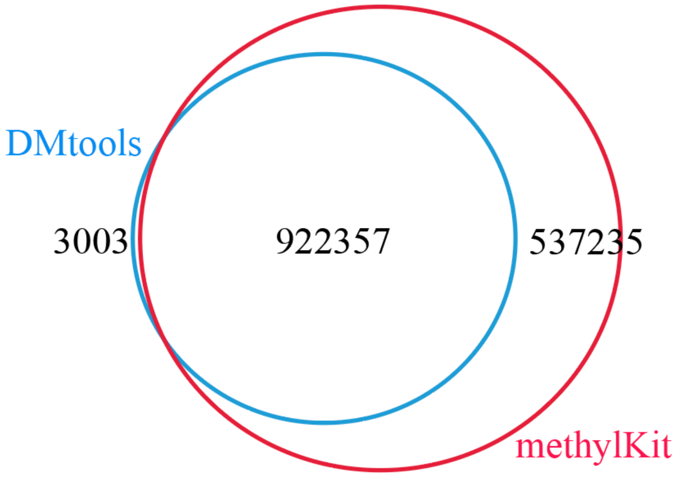


**Fig. S15. The overlap of differential methylation between methylKit and DMtools**

6.4 Absolute difference in DNA methylation level of differential DNA methylation regions

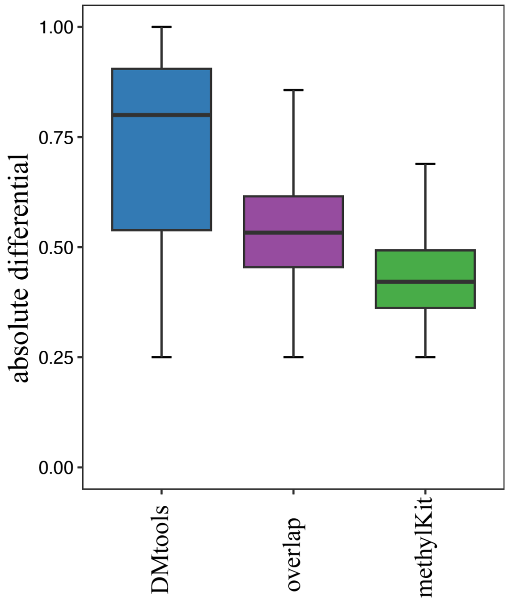


**Fig. S16. The absolute difference in DNA methylation levels of differentially methylated results detected by methylKit and DMtools**

The absolute difference in DNA methylation levels of differentially methylated results specific to DMtools is higher than the overlap portion, while those specific to methylKit are generally lower than the overlap portion.
